# Population inter-connectivity over the past 120,000 years explains distribution and diversity of Central African hunter-gatherers

**DOI:** 10.1101/2021.06.21.449204

**Authors:** Cecilia Padilla-Iglesias, Lane M. Atmore, Jesús Olivero, Karen Lupo, Andrea Manica, Epifanía Arango Isaza, Lucio Vinicius, Andrea Bamberg Migliano

## Abstract

The evolutionary history of African hunter-gatherers holds key insights into modern human diversity. Here we combine ethnographic and genetic data on Central African hunter-gatherers (CAHG) to show that their current distribution and density is explained by ecology rather than by a displacement to marginal habitats due to recent farming expansions, as commonly assumed. We also predict hunter-gatherer presence across Central Africa over the past 120,000 years using paleoclimatic reconstructions, which were statistically validated by dated archaeological sites. Finally, we show that genomic estimates of separation times between CAHG groups match our ecological estimates of periods favouring population splits, and that recoveries of connectivity would have facilitated subsequent gene-flow. Our results reveal that CAHG stem from a deep history of partially connected populations. This form of sociality allowed the coexistence of relatively large effective population sizes and local differentiation, with important implications for the evolution of genetic and cultural diversity in *Homo sapiens*.

## Introduction

The evolutionary history of African hunter-gatherers may hold key insights into patterns and processes behind the evolution of modern human diversity. Recent genomic studies have revealed that these populations represent the oldest and most diverse human genetic lineages, and have been genetically differentiated from one another since the origin of humans (1–3) (Table S5). Therefore, a first question is whether their current ecological niches were also characteristic of early *Homo sapiens* populations. However, genetic data alone cannot determine the geographic distribution of hunter-gatherers in the past nor demonstrate a deep history of adaptation of hunter-gatherers to the environments where they live today. In fact, various studies have proposed that farming expansions within the last 5,000 years (in particular by ancestors of Bantu speakers) would have only recently displaced hunter-gatherers to marginalised regions less favourable to agriculture (such as rainforests and deserts) (4–7).

For example, the central part of Africa between latitudes 5°N and 5°S currently is inhabited by approximately 20 scattered hunter-gatherer ethnic groups (8). These Central African Hunter-Gatherers (CAHG) form a genetic clade that is thought to have diverged from other African populations as far back as 120,000-200,000 years ago (2,9). The lack of any major linguistic specificity between them is often implied to reflect extensive contacts with surrounding farmer populations (8–10), and seen as evidence of recent displacement into marginal forest environments by expanding farming populations. However, anthropologists have remarked on the huge variability in lifestyle, habitat, techniques and tools (11), suggestive of long-term cultural diversification and adaptation to forest environments. Research on the drivers of demography and adaptation of CAHG populations remains extremely limited, which can be partially attributed to the lack of archaeological and osteological data resulting from a rapid disintegration of fossil remains in the rainforest’s acidic soils in addition to social instability in the region (12). Therefore, we are still left with crucial questions regarding the time depth of occupation of Central Africa by hunter-gatherers, the breadth of the niche exploited by earlier populations in the region, and variations in levels of interconnectivity between them at different points in time.

To address those questions, we first compiled ethnographic data on the distribution of 749 camps from 11 hunter-gatherer groups extending from West to East Central Africa. We used them as inputs for environmental niche models (ENMs) to determine the relative influence of several bioclimatic and ecological factors, as well as the presence of farming populations, on the distribution and abundance of Central African Hunter-Gatherers (CAHG) (13–14). We further assess the relationship between the Bantu expansion and CAHG demography by comparing the historical effective population sizes of 9 CAHG populations (N=265 individuals) with those of 15 Bantu-speaking communities (N=677 individuals) that have settled in the region from 20 to 150 generations ago (~600-4,500 years). Then, we use high-resolution paleoclimatic reconstructions and topographic maps to make continuous predictions about where CAHG could have lived over the past 120,000 years and the potential size of their interaction networks. Next, we compile all reliably dated archaeological assemblages ascribed to hunter-gatherer groups in the Congo Basin (N=168) and confirm the models’ ability to predict the location and date of the sites. Last, we contextualise genomic estimates of past demographic trends and population split times with changes in population densities and inter-population connectivity predicted by our model. Our study therefore provides a causal link between past environmental changes and the human population dynamics over evolutionary time, by predicting where and when populations across Central Africa could have exchanged genetic and/or cultural information throughout evolutionary history.

## Results

### Current CAHG distribution reflects ecological adaptation rather than Bantu expansion

We first asked whether the current distribution and density of Central African hunter-gatherers (CAHG) is a product of long-term adaptation to life in the rainforest or instead a recent product of the Bantu Expansion. We compiled ethnographic data on the geographical location of 749 camps from 11 CAHG populations, and then applied the Maximum Entropy (*MaxEnt*) machine-learning algorithm (4) to determine the relative influence of several bioclimatic and ecological factors on the distribution of CAHG (13–14) (Fig. 1A, 1B)(see *Materials and Methods* for alternative model fitting algorithms). ENM performance in the hold-out data was extremely high (TSS=0.55 and AUC=0.87), correctly classifying presences and absences 81% and 74% of the time respectively. The main ecological factor rendering particular areas unsuitable for hunter-gatherer presence was precipitation seasonality (*BIO15*) followed by the annual temperature range (*BIO7*)(Table S8; Fig.S7), confirming what has been proposed for other African regions (15–16). Including predictors relating to Bantu farming populations (*Rural population density* or *Distance to populated places*) neither improved model fit (TSS=0.51 and AUC=0.85 for both models) nor significantly altered the relative contribution of the other variables.

**Figure 1.**
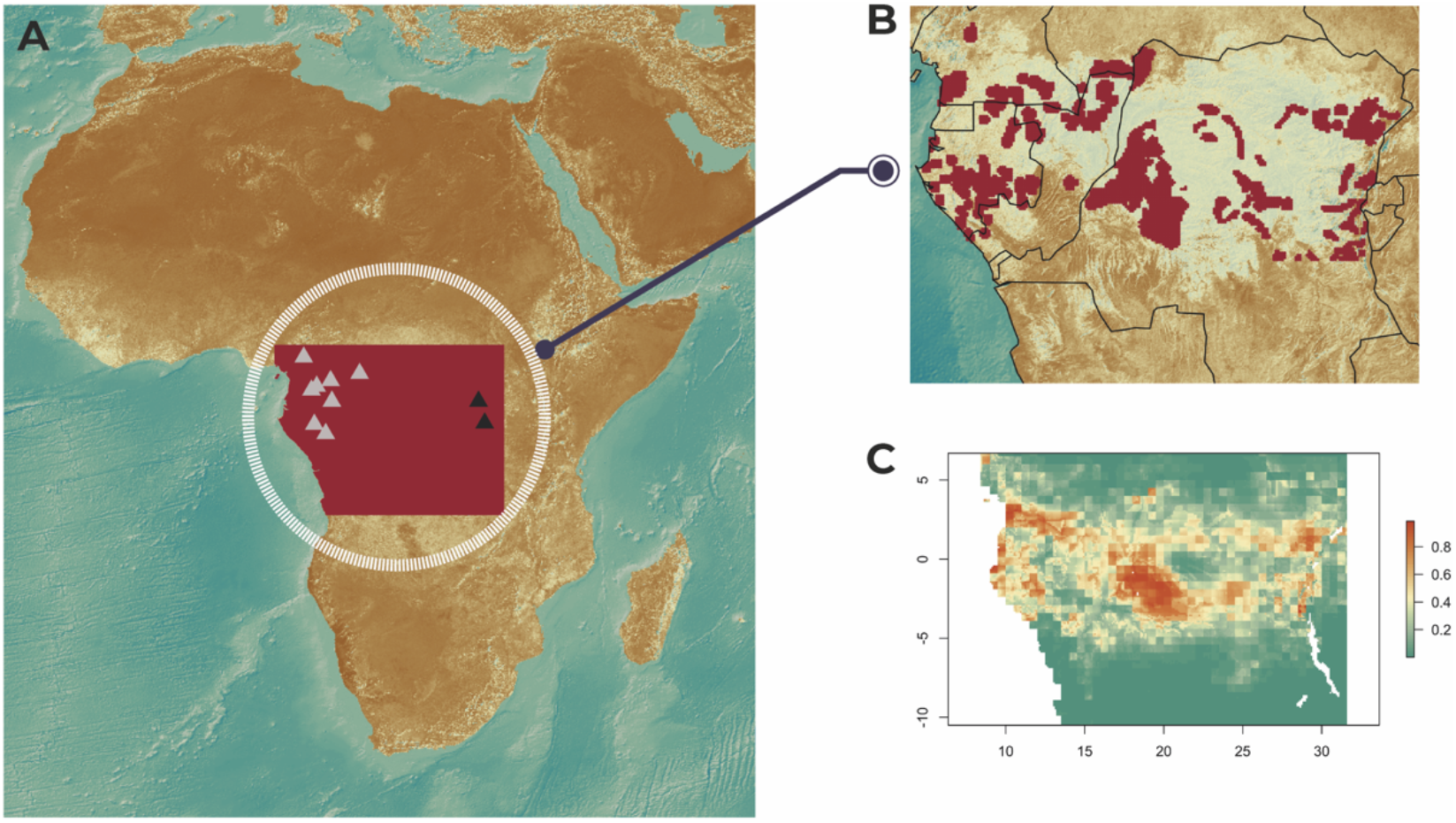
Geographical location of Central African hunter-gatherers. (**A**). Dark and light triangles designate respectively the sources of genomic samples from Eastern (Batwa and Mbuti) and Western CAHG (Baka from Cameroon and Gabon, Bakoya, Biaka, Bedzan, Southern and Eastern Babongo). (**B**) Locations of hunter-gatherer camps used in our bioclimatic environmental niche model (see Materials and Methods).(**C**) Estimated present niche of CAHG using the *MaxEnt* modelling algorithm.

Next, we tested whether the same ecological factors could also predict population density (17). For the 50 map cells where camp size data were available (N=75 camps), we found a positive linear relationship between environmental suitability (estimated by *MaxEnt*) and population density (b= 43.379, t=2.238, *p*=0.030; Fig. S5). We used this relationship to estimate the current CAHG metapopulation size at 191,118 individuals, in close agreement with reported values between 160,000 (11) and 204,000 (18) from census data. In line with previous research, environmental suitability was an even better predictor of the upper limit of population density, as it may be a satisfactory proxy for an absolute constraint on carrying capacity by environmental features (13) (Fig. S5). The slopes of suitability from linear quantile regressions were always significant above the 75^th^ percentile of density, with slopes and R^2^ values increasing with the percentile (Table S10). To test whether those ecological drivers were specific to CAHGs, we applied General Additive Models to predict ecological drivers of farming population densities (SM Text, Fig S8, Table S11). Results showed that bioclimatic variables and biome types affect the two populations differently. For example, high rural population density is associated with temperate forests and large temperature annual ranges, in contrast to hunter-gatherer prevalence which is associated with more stable tropical climates. Thus, our results confirm that the ecological niche of CAHGs and farmers are distinct and that the current range, distribution and populations sizes of CAHG can be predicted by ecological conditions alone (7,19).

### Bantu expansion did not lead to collapse of CAHG populations

Despite several genomic studies on the deep history and demography of African populations (1-2,9,20), their resolution regarding recent times (<10,000 years) is limited. This period is crucial for encompassing the expansion of farming populations, which according to several studies would have caused an opposite contraction in hunter-gatherer populations (7,9,21). We estimated recent changes in effective population size (*N_e_*) from genome-wide SNP data in 9 CAHG populations (265 individuals; Fig.1A) and 15 Bantu-speaking populations (677 individuals; Table S3) from different regions within our area of study (22). To control for the effect for recent admixture with Bantu farmers on the demographic histories of some CAHG populations (4) (Table S6), we estimated ancestry-specific *N_e_* (*ASIBDNe*) after separating CAHG and Bantu ancestry components in hunter-gatherer populations showing significant admixture (see *Materials and Methods*).

We inferred *N_e_* across several Gabonese Bantu speaking populations to range between 10,000 and 100,000 individuals up to around 55 generations ago (~1600 years), in agreement with a previous study (23). However, we found that the *N_e_* of the ancestors of CAHG in this region and Bantu-speaking peoples from Cameroon and Uganda show similar rather than opposite trends. Interestingly, during this period, we also observed a population decline among the Baka of Cameroon starting from 2,500BP coinciding with a rapid fragmentation of the rainforest in the area occupied by this population (24). Seidensticker et al. (23) inferred a collapse of Bantu-speaking forest dwellers during the subsequent period 1600-1200BP (53-40 generations ago) based on archaeological data and declines in N_e_. When including additional farming groups from our study, we found that whilst some Gabonese Bantu-speaking communities would have suffered demographic bottlenecks during this period (Fig. 2C), this was not the case for Bantu speakers from Cameroon or Uganda and most CAHG (Fig. 2A-B). In fact, between 1200-600BP (40-20 generations ago), we see evidence of increases in the N_e_ of CAHG from the Northwestern and Eastern regions (Although results for the Mbuti should be taken with caution given our limited sample size for this population)(Fig. 2A-B; Fig.S14) yet a slight decrease in the effective size of Gabonese CAHG. This is most likely due to an episode of aridity and deforestation in Gabon spanning 1240-550BP (25).

**Figure 2.**
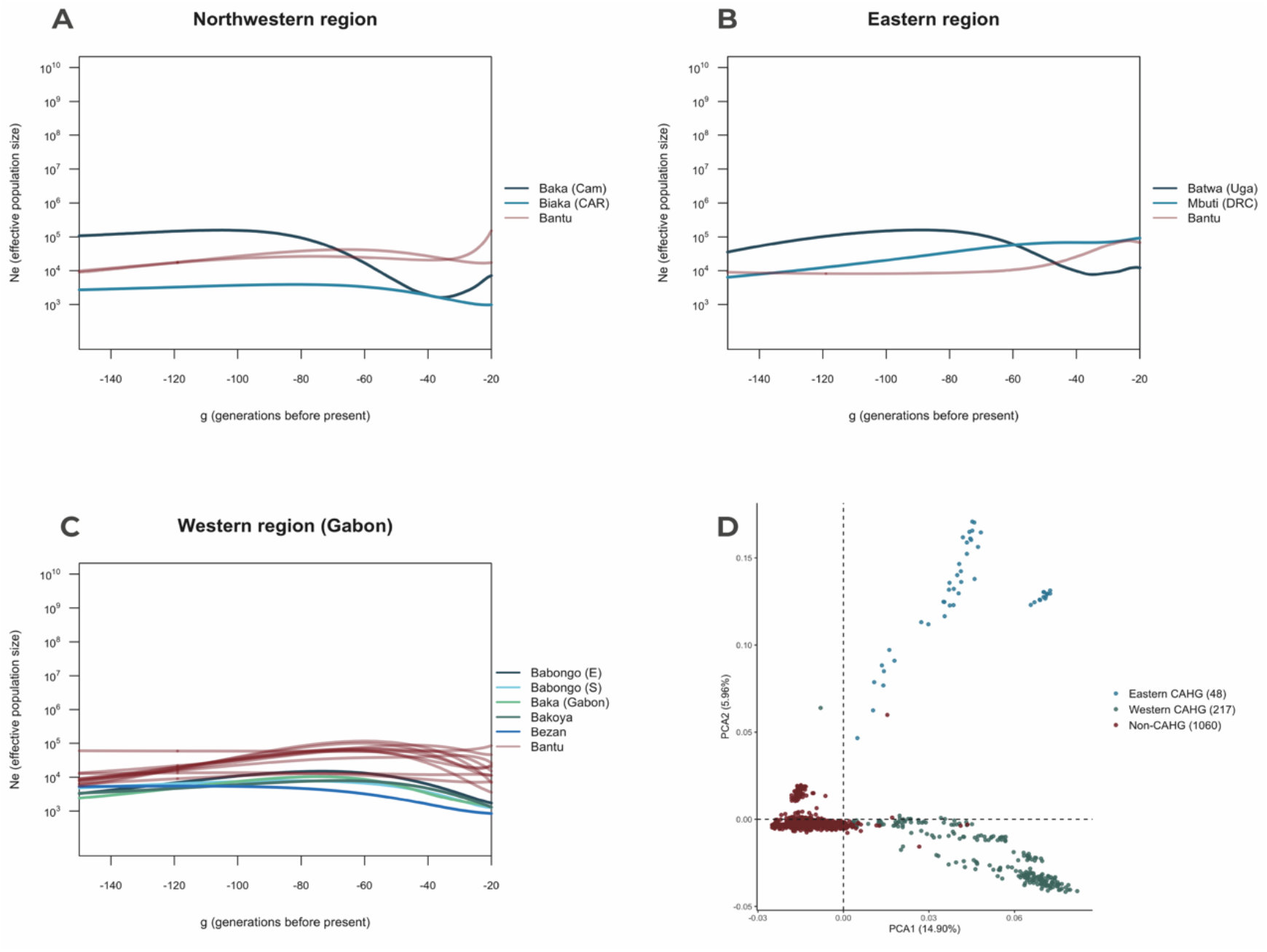
Ancestry-specific effective population size (*N_e_*) estimates for CAHG and Bantu-speaking populations from 4,500 years B.P. until 600 years B.P. and structure of genetic variation. (**A**) *N_e_* estimates for northwestern groups include Baka and Biaka (CAHG) and Bantu-speaking Badwee and Nzime from Cameroon and the Central African Republic. (**B)** N_e_ estimates for Eastern groups Mbuti and Batwa (CAHG) and Bantu-speaking population Bakiga from the Eastern part of the Democratic Republic of Congo and Uganda. (**C**) *N_e_* estimates for Southern Babongo, Eastern Babongo, Baka, Bakoya and Bezan (CAHG) and Bantu-speaking Akele, Benga, Duma, Eshira, Eviya, Fang, Galoa, Makina, Ndumu, Obamba, Orungu and Shake from Gabon. Each generation represents 30 years. (**D)** Principal Component Analysis showing genetic diversity of 40 African populations (N=1325 individuals), including CAHG populations (Mbuti, Biaka, Baka from Cameroon and Gabon, Bezan, Bakoya, Batwa, Southern and Eastern Babongo).

Together, our results suggest that the population dynamics of hunter-gatherers and farmers are independent, and subject to region-specific environmental factors as opposed to competitive exclusion (17). Consequently, CAHG demographic trends reflect a long history of adaptation to Central African ecologies as opposed to a recent occupation following the expansion of agriculturalist populations.

Our ADMIXTURE analyses (26) also indicate a higher proportion of Bantu ancestry in Western (mean=25%) than Eastern CAHG (mean=16%; Fig.S13), consistent with archaeological and linguistic evidence (27–28) proposing a Bantu origin somewhere between Eastern Nigeria and Western Cameroon. Nonetheless, there were many individuals that did not carry signs of Bantu admixture (Fig. S13).

### Ecology predicts viable CAHG populations in tropical forests throughout the last 120,000 years

After establishing that ecological factors are the main determinants of locations of current hunter-gatherers in Central Africa, we then applied our model to predict locations of past CAHG, and to estimate how their level of interconnectivity may have varied across regions. We employed a bias-corrected time series of global terrestrial climate and vegetation (15) to project our ENM model into 1000- or 2000-year time slices from the present up to 120,000 BP (see *Materials and Methods*), obtaining suitability maps in our area of interest for each time slice. Similarly, we used our estimated linear relationship between environmental suitability and grid cell population density to obtain metapopulation size estimates at each time slice. Our results indicated that repeated episodes of abrupt climatic changes and habitat fragmentation in Central Africa since the Last Interglacial (15,29-30) would have caused drastic expansions and reductions of the suitable range for CAHG groups (Fig. 4B; Fig. S9; Movie S1). Whilst this would have resulted in demographic fluctuations, the populations of hunter-gatherer could have remained viable in Central Africa, with a potential census metapopulation size never falling below 160,000 individuals (Fig. S19). This suggests that CAHG would have been able to maintain their niche and relatively stable population sizes throughout evolutionary history. Unlike approaches relying exclusively on genetic or linguistic evidence, our model directly situates geographically and ecologically the locations of putative ancestors of CAHG. Contrary to claims that Central African environments would not support the presence of human groups independent of agriculture (6-7, 18), our results add further support to previous linguistic and genetic studies that modern CAHG are descended from ancestral groups whose adaptations to the rainforest may extend back into the Late Pleistocene (1,8-9,20).

### Ecological predictions match the Central African archaeological record

Amongst other factors, high temperatures and abundant precipitation in rainforest environments limit the preservation of organic remains (12,31). Thus, the archaeological record of the region is limited and biased towards non-forest areas. Therefore, rather than to delimit the niche of CAHGs in the past (32), we use the archaeological record to test our model’s ability to predict the location and dating of known archaeological sites occupied by ancient hunter-gatherers. We compiled all published ^14^C dates (N=168) from hunter-gatherer archaeological sites from the Middle Stone Age onwards (Data S2-S4; Fig.S20; Fig. 3A; *Materials and Methods*). We removed sites with multiple dates within the same 1000-year time slice, resulting in a sample of N=118 dated sites. Analyses showed that archaeological sites were almost twice as likely to be found in cells predicted to be suitable in our model than in randomly selected cells in their corresponding time slices (Observed=51, Expected=28, χ^2^ =12.677, df=1, p-value<0.0001)(Fig. 3B). We then performed 1000 random permutations of the ^14^C dates of our CAHG sites, and confirmed that the number of sites falling in suitable cells when assigned their correct date was higher than predicted by any of the permutations (Fig. 3B), mean across permutations=31.9, SD=5.08; t=106.19, df=999, *p*<0.0001). Furthermore, our model performance was not related to the sites’ dates (r=0.003, t=0.393, df=107, p=0.968). This indicates that the location of archaeological sites is not only determined by geographic or topographic features, but by how suitable such locations would have been for hosting hunter-gatherer populations at different points in time. The combined results of our models and the archaeological site distributions support the idea that CAHG have long been adapted to life in tropical environments, also consistent with the available fossil evidence (Table S12)(33). Moreover, that expansions and contractions of suitable ranges have influenced the demography of CAHGs.

**Figure 3.**
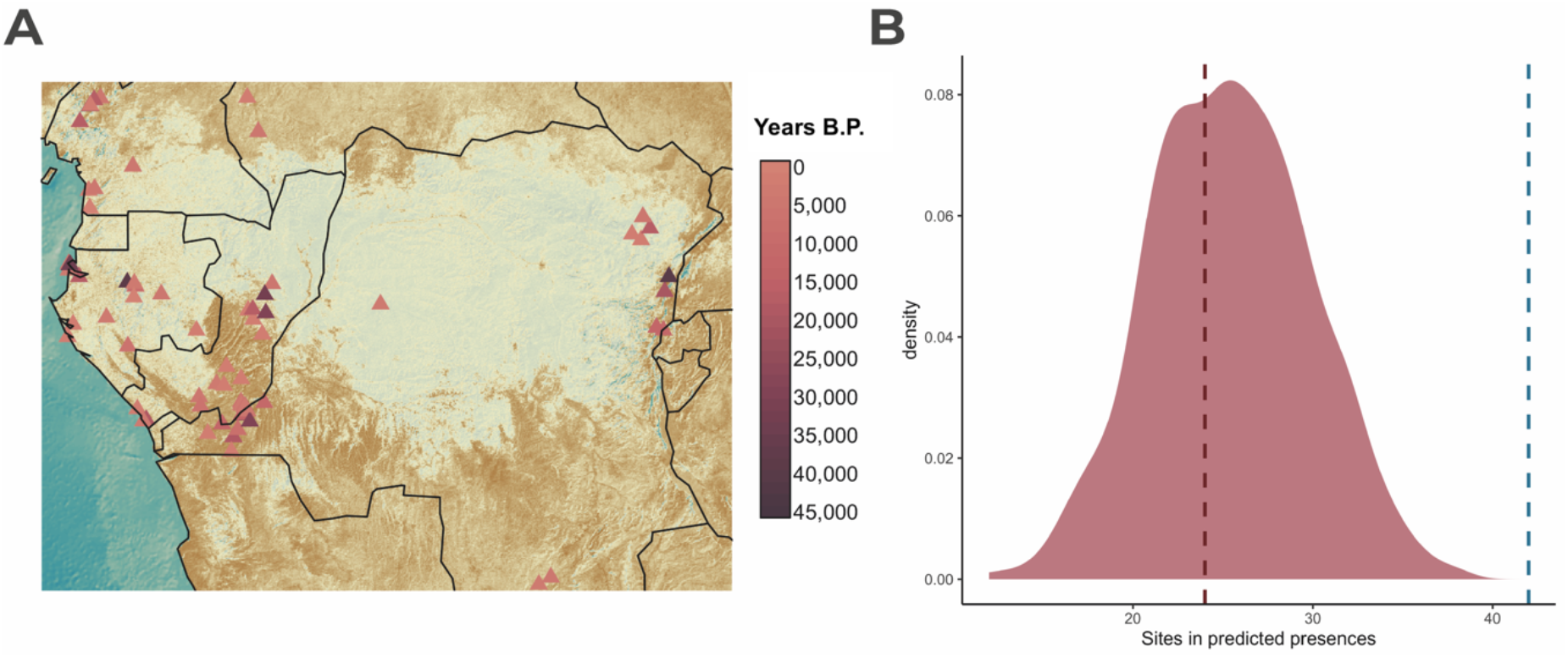
Ecological modelling of archaeological site distribution in Central Africa since 47,000BP. **(A)** 118 Archaeological sites included in our analyses and their ^14^C dates. **(B)** *MaxEnt* predicted the presence of 51 sites in suitable cells (blue line). Distribution shows number of sites in predicted presences across 1000 randomisations of site dates. Dashed pink line indicates-expected cumulative number of sites in predicted presences when randomising their spatial location at each time period.

### Ecology-driven changes in the range size and connectivity of CAHG populations explain population and genetic splits

To understand how environmental changes in Central Africa affected the deep evolutionary histories of African hunter-gatherers and their levels of genetic and cultural exchange, we modelled metapopulation sizes and connectivity over the last 120,000 years. For each time-slice, we plotted all camps predicted by our ENM. To estimate their degree of interconnectivity, we calculated the cost of movement around each camp using Tobler’s Hiking Function (34), which takes into account topographical features, rivers and water masses that represent challenges to mobility. Then we computed the total number of neighbouring camps within a predicted 7-hour walk from it. This radius corresponds to Cavalli-Sforza and Hewlett’s empirically derived (21) average “half range” of Aka Hunter-Gatherers, and matches posterior estimates of lifetime ranges of other CAHG populations (35–36). To evaluate how changes in connectivity affected genetic differentiation between populations, we compiled recent genomic studies of CAHG demographic histories and their predicted split times, and compared our predicted range sizes and interconnectivity levels to genomic estimates of population splits obtained from whole-genome studies using cross-coalescence methods (Table S5; Fig. 4C).

**Figure 4.**
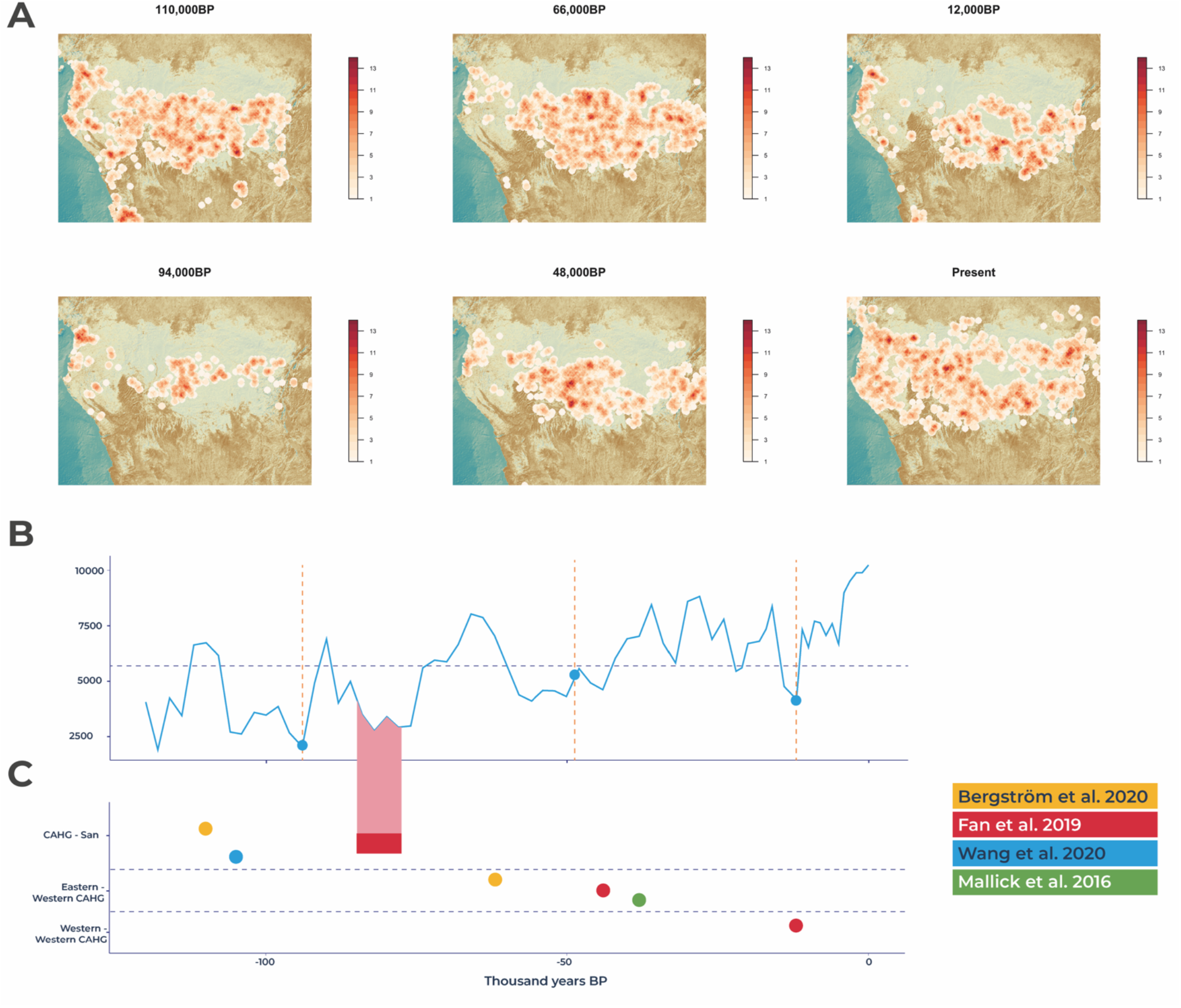
Predicted dates of genetic splits between CAHG populations match decreases in predicted CAHG range sizes and group interconnectivity. **(A)** Predicted connectivity from our *MaxEnt* model at the time periods corresponding to the genetic estimates of population divergence times as well as before them. The predicted number of camps at each time period was randomly distributed across the cells with predicted presences, and the number of other camps within a 7hr walk of each camp calculated. Darker shading indicates a greater number of camps within a 7hr walking distance of one another. (**B**) Range size of CAHG across time predicted by our *MaxEnt* model. Vertical dashed lines indicate estimates of genetic split times by averaging all available studies for a particular event. Horizontal dashed line indicates average size of suitable range for CAHG. **(C)** Estimates of divergence times between African hunter-gatherer populations using cross-coalescence methods on whole-genomes. Coloured segments represent midpoint of cross-coalescence rates.

Our data show that at 94,000 BP, the midpoint of the divergence between the CAHG clade and the San (the earliest human genetic split) estimated across different studies (1,37), coincides with our model estimation for the culmination of a drastic reduction in the suitable range for hunter-gatherers as well as decreased connectivity (Fig. 4A-B). The few hunter-gatherer populations in Central Africa would have been small and relatively isolated, compatible with separation between San and CAHGs (Fig. 4B).

Similarly, the midpoint of separation times between Eastern and Western CAHG at 48,000BP falls in a period during which our model predicts decreased connectivity between East and West parts of Central Africa lasting from 66,000-42,000BP (1,37-38). Finally, at 12,000 BP (the predicted genetic split between different Western CAHGs)(1), our model shows the fewest number of suitable cells during the past 50,000 years. This is particularly pronounced in the West, where suitable ranges get restricted to those in the coast of what is now Cameroon. In summary, our data show that genetic estimates of splits among CAHGs mostly take place during periods when the size of the suitable range falls below average and connectivity between regions is reduced.

### Evidence of gene flow between Eastern and Western hunter-gatherers in the last 2,500 years

Besides periods of ecological niche contraction favouring population splits, our model also predicts periods of noticeable niche expansion and gene flow between previously separated hunter-gatherer groups. We thus used patterns of sharing of Identical-by-descent genetic segments (IBD) between populations to provide insights into recent contact among populations with deep common ancestries. We analyzed IBD blocks in three categories: 1–5, 5–10, and >10 cM (Fig. 5), roughly corresponding to time intervals of 2500–1500 years ago, 1500–500 years ago, and 500–0 years ago respectively (39–40). During the three-time intervals, analyses identified widespread gene flow among Western groups, as well as between Eastern and Western CAHG (Fig. 5; Table S13). High levels of connectivity in the past 2500 years between geographically and genetically differentiated hunter-gatherer groups argue against population decrease and fragmentation after Bantu expansion. Although our data does not allow dating inter-group connectivity at previous times, we also find evidence of distinct sharing of private alleles between all pairs of CAHG populations (Fig. S18), again indicating gene-flow across Central Africa (41). Together with our simulations, this strongly suggests that the histories of fragmentation and connectivity among CAHG are equally deep. This pattern is compatible with recent studies indicating that rather than clean splits, genetic separations within Africa were gradual and shaped by ongoing gene flow (20), and with evidence of gene flow between Mbuti (East), Biaka (West) and San (South) until at least 50,000 years ago, and between Biaka and Mbuti (East) until the present day (37,41). If this scenario is correct, the evolutionary history of *Homo sapiens* must be seen in the light of such long-term demographic dynamics involving both population fragmentation and interconnectivity.

**Figure 5.**
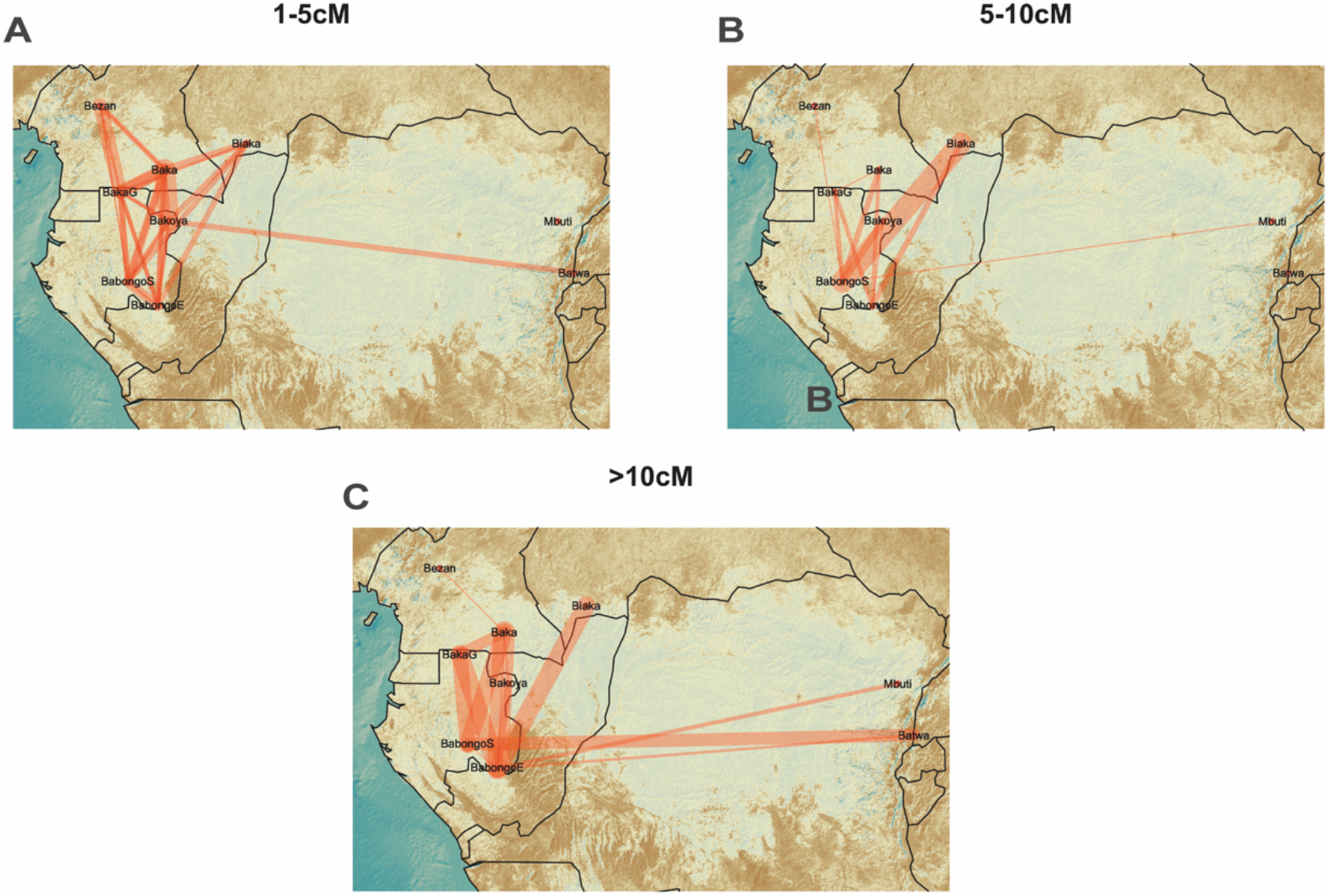
Recent genetic connectivity between CAHG populations. Network visualizations of the average number of IBD segments shared per cross-population individual pairs with identified IBD blocks in the range of: **(A)** 1–5 cM (2500–1500 years ago), **(B)** 5–10 cM (1500–500 years ago) **(C)** and over 10 cM (500-0 years ago). Thicker lines indicate levels of gene flow as identified by a higher probability of sharing IBD blocks of the specified length.

Our ecological model can therefore recover the timing of major genetic events of separation both within central Africa, between central African and other hunter-gatherers, and may account for the relatively high levels of genetic differentiation (F_ST_) among CAHG groups when compared to nonhunter-gatherer populations inhabiting the same areas (ANOVA: *F*-value=631.6, df=2, *p*<0.0001) (42)(Fig.S11). Thus, our prediction of multiple cycles of isolation and reconnections over the last 120,000 years would account both for the maintenance of high effective population sizes throughout the Late Pleistocene (~30,000 individuals) in spite of the small band sizes documented in ethnographic records (1,20,37,43-45), and of relatively high levels of genetic diversity compared to the more recent descendants of Bantu-speaking farmers.

## Discussion

We have modelled the role of environmental change since the last interglacial on the distribution, density, and dynamics of hunter-gatherers in Central Africa, a key geographical bridge between Northern, Western, Eastern and Southern Africa but nonetheless traditionally left out from human evolutionary studies (12). Contrary to common assumptions, the current range and abundance of CAHG is mainly determined by long-term adaptation to ecological conditions rather than recent displacement by expanding Bantu-speaking farmers (28,46-47). Despite vast fluctuations in environment and available niche during the Late Pleistocene, CAHGs would have always maintained relatively large, viable and distinct populations (15,29-30). The fact that our modelled relationship between ecology and CAHG presence is consistent with the location of past archaeological assemblages and human remains suggests that as well as having remained genetically distinct for a very long time, contemporary African hunter-gatherers also occupy similar habitats to those exploited by their ancestors. It also supports recent morphological and genetic evidence that early hunter-gatherer populations in Africa were highly structured into semi-rather than fully isolated groups (1,9,20,48). This form of sociality characterised by a combination of local differentiation and partial connectivity, facilitated by the fluid structure of hunter-gatherer bands, may also explain the ability of early hunter-gatherer groups to maintain relatively large and stable effective population sizes (49–51) despite regular episodes of environmental fragmentation.

Although the increasing availability of paleoenvironmental reconstructions has resulted in a consensus over the extreme climatic variability characterising African environments during human evolution (29–30), few studies had attempted to directly test the effect of changing ecologies on human population dynamics (19,52-53). We hope that future research adopts similar spatio-temporally, ethnographically explicit approaches to further shed light on both the origins of modern human ancestry and the flexibility and adaptive potential of the human foraging niche. By determining the role of changing ecosystems in shaping variation within and between groups of humans across Africa, our study helps to characterise the breadth of the ecological niche exploited by early *Homo sapiens* populations known to extend well beyond coastal and savannah environments, and may provide valuable insights into past between-group relationships, as well as their relative contribution to present genetic, behavioural and cultural diversity.

## Materials and Methods

### Hunter-Gatherer presence data

Our study area extended 6.7°N, 10.5°S, 31.6°E, 8.4°W, comprising five Central African countries (Cameroon, Central African Republic, Gabon, Republic of Congo and the Democratic Republic of Congo). Within it, we obtained georeferenced location data for a total of 749 documented CAHG camps from a combination of primary literature and a previous study by Olivero et al.(13). We excluded camps that are the result of forced relocation after government re-settlement programs (54) (S1 Data).

Following Olivero et al. (13) we generated a buffer zone of 20km of land around each camp representing a theoretical unit area of land liable to be exploited for natural resources. To determine this area, Olivero et al. (13) first determined the mean radius (18.5 ± 1.0 km) encircling a camp, by using average subsistence area. They also obtained the average travel distance covered for subsistence activities from 36 studies (21.0 ± 3.65 km; (35, 55-57). From these two measures, the buffer zone around each camp was chosen to be of 20km.

We then applied this buffer to all camps in our database to plot onto a 0.1° × 0.1° map of the study area. This resulted in 4,577 grid cells of CAHG presence, out of the total of 36,277 cells that covered the entire study area (Fig.1B). We then considered absences or background (depending on the modelling algorithm used) to be those grid cells not included in the presence-grid-cells (see below).

### Environmental and paleoenvironmental data

We made use of the same newly published high-resolution (0.5°) bias-corrected time series of global terrestrial climate and vegetation data covering the last 120,000 years (15) for both building our models and extrapolating them into the past. Gridded reconstructions are available at 2,000-year time steps between 120,000 and 22,000 BP, and 1,000-year time steps between 21,000 BP and the modern era. The data include 17 bioclimatic variables, which have been used extensively in environmental niche models as well as reconstructions of global biomes, leaf area index and net primary productivity. For interpretability purposes, we aggregated the 27 biomes simulated in Beyer et al. (15) into the same megabiomes as described in the original paper. In addition to bioclimatic variables, we obtained topographical and hydrographical variables from the US Geological Survey (58) as well as from HydroSHEDS (Table S1).

After obtaining a selection of climatic, bioclimatic and vegetation variables that are thought to affect the distribution of forager populations from previous literature, to avoid multi-collinearity issues, we checked for pairwise correlations among them (Table S1; Fig.S1). Following Dormann et al.(59), we used an adapted version of the *select07* method to identify all pairs of variables with |*r*|<0.7 and remove the less important variable in terms of explained deviance. The final models included 9 predictors (Fig.S2; Tables S8-S9).

To fit the set of models (see below) designed to test the relative influence of the presence of farming populations whose ancestry can be traced to the Bantu expansion on the current distribution of CAHG, we also obtained data on the *Rural population density* at each cell in our map as well as its as *Distance to populated places*. The former was taken from the Relational World Database II (RWDB2) and the later from the LandScan 2008 High Resolution Global Population Data Set (60), excluding any areas less than 2-km far from urban areas.

### Archaeological data

We compiled all published calibrated ^14^C dates from archaeological sites in the area of Central Africa covered by our study dating from the Middle Stone Age onwards. Their dates range from the present to 46,000, corresponding to the limit of radiocarbon dating (23-24,61). First, we removed ^14^C dates based either on carbonates-possibly affected by carbon reservoir effects-or with associated errors greater than 1000 years (the resolution of our paleoclimatic) as well as dates associated with dubious stratigraphy or missing laboratory codes. (Data S3). The final sample after this stage comprised N=930 sites (Data S3). However, for the present study, only sites occupied by hunter-gatherers are relevant, hence, we needed to remove any sites that could have been occupied by Bantu populations.

From the primary literature, we obtained the chrono-cultural affiliation of each of the sites as well as checked the description of the assemblages to make sure the criteria coincided. Presence of farming populations in archaeological sites was inferred from the occurrence of *Pennisetum glaucum* (Pearl millet), *Elaeis guineensis* (Oil palm), iron metallurgy or pit features (23–24). Those sites that did not have such features but that had been assigned to either the “Neolithic”, “Early Iron Age” or “Recent Iron Age” by their original authors were also considered to represent farming populations and hence excluded from our analyses. In addition, we checked the agreement of our chronocultural affiliation and assessment of dating reliability with that reported in recent summaries of the archaeological findings in Central Africa by Garcin et al.(24); Oslisly et al.(62); Morin-Rivat et al.(63); Cornelissen (31) and Seidensticker et al. (23). The remaining dates, that is, those with a chronocultural affiliation within the Stone Age (Middle or Later), no evidence of farming population occupation and reliable ^14^C dates (according to the criteria outlined above) were assumed to represent hunter-gatherer sites in the past. We classified these dates into two categories, the first one including the whole sample (N=168 dates) and the second including the subset of sites with associated lithics from hunter-gatherer populations (N=151 dates)(Data S2; Fig. S10). We did not re-calibrate the dates for the present study.

This sample is likely to be overly conservative, as our selection criteria implies that any sites that would have been co-occupied by hunter-gatherers and farming populations, where trade between foraging and farming groups would have traded tools or where hunter-gatherers would have utilised or cultivated any of the above-mentioned plants (something that is very common among present populations; e.g. (64–65) would have automatically been removed from the sample. However, given that our aim is to test the strength of the evidence for an ancient occupation of Central Africa by hunter-gatherers, a conservative approach is desired. To further verify the robusticity of our results to different site-selection criteria, we ran the analyses on two additional datasets: one excluding several disputed dates and another one excluding sites attributed to hunter-gatherers but with evidence of pottery use (66)(Data S2; Fig. S22-S23).

### Genetic data

Illumina array data were accessed from three published datasets (21,67-68). These data were filtered for relatedness >0.0886 with KING (69). A random individual was chosen from each related pair to give a total of 1325 individuals (Table S2). The data were then pruned for MAF <0.05 with PLINK (70). A pruned version of the dataset was created with PLINK --indep-pairwise 50 5 0.2 to account for linkage disequilibrium. The non-pruned dataset comprised 1325 individuals and 555,630 SNPs and the pruned dataset comprised 1325 individuals and 213,927 SNPs. The unpruned dataset was used to calculate runs of homozygosity and heterozygosity using PLINK. The pruned dataset was used to assess the structure of genetic diversity within our area of study by calculating Wright’s F_ST_ statistics between all pairs of populations (42) using PLINK and for decomposing genetic variants into principal components using smartPCA with 10 outlier iterations (71).

### Environmental Niche Modelling of distribution of CAHG

Environmental Niche Models (ENM) or Species distribution models (SDMs) are an extremely powerful tool from ecology (72). They mathematically relate occurrences of a particular species (in this case “species” corresponds to georeferenced hunter-gatherer camps) and the bioclimatic or ecological features of the areas it inhabits to produce a model that by identifying the “realised niche” of the species (73) can predict its potential geographic distribution based on suitable environmental conditions. Increasingly, ENMs have started being used to also understand the non-random distribution of human populations or cultural traditions in the present and in the past (13-14,32).

To model the potential distribution of CAHG throughout Central Africa, we used Maximum Entropy (MaxEnt)(72), one of the most widespread and best-performing techniques for ENM. Nonetheless, to minimise the potential of our model predictions to be reliant on the particular model algorithm used, we also performed the models using the Favourability function obtained from Generalized Linear Models (GLMs), and compared their output with that obtained from MaxEnt (SM Text). MaxEnt is a type of non-parametric machine-learning (ML) method. MaxEnt takes a list of species presence locations as input, as well as a set of environmental predictors across a landscape divided into grid cells. From this landscape, MaxEnt extracts a sample of background locations that it contrasts against the presence locations. The model then estimates a Relative Occurrence Rate (ROR) at each grid cell, as a function of the environmental predictors at that location (74). The ROR is the relative probability that a cell is contained in a collection of presence samples. MaxEnt allows fitting very complex, highly non-linear response shapes (72). However, it also limits model complexity – and, hence, protects against overfitting by regularization: a penalty for each term included in the model and for higher weights given to a term (74). This is important given our primary aim to use the model to make predictions in environments that differ to those used to fit (in our case, paleoenvironments). We used all feature classes (linear, hinge, quadratic, product, threshold and discrete), as our predictors are a mixture of categorical and continuous variables.

A key objective of our study was to use the fitted species distribution models to project hunter-gatherer distributions under past climate scenarios. MaxEnt is the preferred model for extrapolating species’ distributions to new environments because it is “clamped,” that is, it extrapolates in a horizontal line from the most extreme environmental values in the training data set (72). Previous studies have also used this algorithm to model palaeolithic niches of human populations (32). Predicted suitabilities obtained from from MaxEnt were converted to presence/absence predictions using the optimum threshold value maximizing the sum of sensitivity and specificity (75).

The ability of the model to predict hunter-gatherer camp occurrence was assessed using standard cross-validation procedure (80% random sample for calibration and 20% for validation; 500 repeats)(76). The predictive power of the binary models was determined by testing the accuracy of predictions made for the validation dataset (not used to build the model) by calculating the area under the curve of a receiver operating characteristic plot (AUC), Cohen’s Kappa Statistic (77) and the True Skill Statistic (TSS) that unlike the Kappa statistic is independent of prevalence and therefore in recent years has been considered a more appropriate way of measuring the performance of ENM (78).

Last, to assess the robusticity of our model predictions to potential slight changes in the flow or position of minor rivers, we fitted a second set of models excluding “distance from water masses” as predictor.

### Estimation of past distributions of CAHGs

The logistic output of MaxEnt consists of a grid map with each cell having an index of suitability between 0 and 1. Low values indicate that conditions are environmentally unsuitable for the presence of hunter-gatherer camps, whereas high values indicate that conditions are suitable.

After calibrating our model using the current range of hunter-gatherers (i.e. the present locations of hunter-gatherer camps), we projected it into the each 1000- or 2000-year time slice from the present to 120,000BP to obtain suitability maps in our area of interest for each time period.

### Estimation of camp and CAHG population density in the present and past

The relationship between abundance and distribution range has been extensively studied in biogeography e.g. (79–80) Abundance may be determined by limiting physiological variables or the ecological characteristics of species (17), which do not always exhibit regular spatial patterns (79). If these limiting factors are the same as those that also condition species presence, then models accounting for species occurrence could be useful in providing information on species abundance. Hence, from our distribution models, we estimated past and present population densities.

Since previous studies have found a positive association between population *density* and environmental suitability), the relationship between hunter-gatherer population density and environmental suitability was examined in 64 grid cells (N = 100 camps) for which camp-size data were available. We calculated CAHG population densities from the sum of all CAHG population figures reported for the same 0.1° × 0.1° grid cell (123 km^2^ at the Equator). We examined the shape of the population—suitability values point cloud, after population-size outliers were eliminated following Tukey (81) i.e., if population size > Q3 + 1.5 × (Q3—Q1), where Q1 and Q3 are the first and the third quartiles, respectively] and found a typical wedge-shaped relationship.

We then used ordinary linear regression to test the significance of a positive relation between the population size at particular grid cells and their suitability values. In addition, since previous work suggests data on species distribution is particularly useful to predict the upper limit of species abundance (see Olivero et al. (13) for evidence of the same relationship when considering CAHG camps) we also fitted linear quantile regressions to the 50^th^, 55^th^, 60^th^, 65^th^, 70^th^, 75^th^, 80^th^, 90^th^, 95^th^ and 99^th^ percentiles, and the R1 measure (weighted sum of absolute residuals) was calculated in each percentile as a local measure of goodness-of-fit (82).

We use the term metapopulation here to encompass all spatially separated populations of CAHG groups. (Note that Olivero et al. (13) define it as all spatially separated CAHG groups that may interact to some extent). To estimate metapopulation size, first, we calculated the average CAHG population size empirically observed in grid cells for which population sizes were available (after removing outliers. Using the coefficient of the slope from the linear regression between suitability and population density, we then calculated the potential population size (PPS) for every grid cell in the study area, according to their suitability values. Finally, we summed all PPS values for the entire study area, but applied the following correction to take territoriality into account:

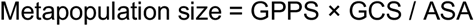

where the metapopulation size is the net potential population size; GPPS is the gross potential population size resulting from the sum of the PPS values; GCS is the size of a grid cell (i.e. 123 km2); and ASA is the average subsistence area estimated for CAHG (i.e. 1,079 km^2^).

### Estimating the degree of connectivity of hunter-gatherer populations

One of the largest challenges for estimating the potential for contact, and therefore the effective population size exchanging genetic and cultural knowledge between past and present hunter-gatherer populations is the very limited availability of data on hunter-gatherer mobility, movement ecology and resource use.

As a first estimate of the degree of connectivity of the predicted hunter-gatherer camps at every time step, we will use Cavalli-Sforza and Hewlett’s (83) estimation of the average “half range” of Aka Hunter-Gatherers, defined as the median of the distance from their current place of residence to the places they had visited at least once during their lifetime. This measure was used to estimate a maximum distance between camps whose members could have interacted with one another. Their empirically derived measure was of 34.5km or around 7 hours of travelling time around a camp, is congruent with posterior estimates of life-time ranges of other CAHG populations (43–44). Nonetheless, it is important to note that among Aka CAHG, rare movements between 60–80km have also been recorded (83) mostly to visit family members living far away or to obtain work in villages (56).

Since the number of cells with predicted hunter-gatherer presences from our model (at any time period) would be larger than the actual number of hunter-gatherer camps (since all cells within a 20km radius of a camp are considered to contain “presences” – as this represents the average subsistence area used by camps), we derived the ratio of camps to cells with presences from the original map, which was found to be of 0.16. Hence, for every time period, the predicted number of cells with presences was multiplied by 0.16 to obtain the predicted number of camps at that timeslice. Then, to estimate the distribution of camps from which to calculate connectivity, for each timeslice, the predicted number of camps by each of the algorithms was randomly distributed across the cells with predicted presences.

However, our study area is a mosaic topographical features, rivers and water masses that would have presented logistical challenges to hunter-gatherers. To take such logistical challenges into account when considering camps’ potential connectivity, we used the GTOPO30 Digital Elevation Map (58) to calculate the anisotropic accumulated cost of movement around each camp. We will do this with Tobler’s (34) hiking function that estimates speed, taking into account the slope and its direction, and is the most popular cost function in archaeological least-cost path calculations (84-85). Then, to estimate the degree of connectivity of each camp, we calculated the total number of other camps within 7 hours of walk from that camp.

### Validation of the model using archaeological assemblages

A great limitation for assessing our the adaptability of the ancestors of CAHG to rainforest environments is that high annual temperatures and abundant precipitation generate acidic conditions in the soil, which limit the prservation of organic remains such as pollen, soft plant tissues, bone, and wood (12,31) (Fig. S20). This means that the archaeological record of our area of interest is not only limited but biased toward non-forest areas where preservation is better. Consequently, following approaches that rely on the location of the archaeological sites themselves to delimit the niche of hunter-gatherers would not be justified (e.g. d’Errico et al.(32)). Instead, we can use the archaeological record to test whether our modelled relationship between ecological variables and CAHG presence in the present (using an exhaustive dataset) also predicts the instances of CAHG presence in the past. In other words, we used the existing archaeological record to assess whether the continuity of the niche occupied by CAHG through time.

To test whether our estimated suitability landscapes in the past matched areas that were actually occupied by hunter-gatherer populations, we obtained the suitability values of the grid cells where archaeological sites are located at the time corresponding to their ^14^C date (rounded to the nearest 1000-year or 2000-year according to the resolution of our paleoclimatic reconstructions for each time snap). We used the optimum threshold obtained when building the model for MaxEnt-derived predictions (0.31) to binarize those values, and in so, obtain the number of archaeological sites located in cells with predicted presences. To avoid sampling biases from sites with multiple dates close to one another, only one data point per 0.5×0.5° cell per 1000- or 2000-year interval was included (Data S4).

We then compared these values with the number of sites we would expect to observe in cells with predicted presences if sites were distributed randomly across the suitability landscape at each time period (that is, if suitability and archaeological site presence were independent of one another). In this way, we were able to calculate whether the likelihood of finding archaeological sites in cells with predicted presences was above chance.

To calculate the expected number of sites in predicted presences, we took, for each time period:

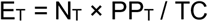

Where ET is the expected number of sites in predicted presences at time period *T*, N_T_ is the number of sites with ^14^C dates within time period T, PP_T_ is the number of predicted cells with presences at time period T, and TC is the total number of grid cells in our map (36,277).

We added the expected number of cells with predicted presences across all time periods and divided it by the total number of archaeological sites to obtain the proportion of archaeological sites that would lay on cells with predicted presences if suitability and site location were independent of one another. We compared this with the observed proportion of sites located in cells with predicted presences.

We further validated the projections of our model by testing whether changes in the suitability landscapes over time indeed determined the spatial locations occupied by CAHG. To do this, we performed 1000 random permutations of the ^14^C dates of our sites. At each permutation, each site was randomly assigned one of ^14^C dates from the list of dates from our sites, and the suitability value at the location of the site, at the time-period corresponding to the assigned ^14^C date extracted. Then, for each permutation the total number of sites falling on predicted presences was counted, in so obtaining the number of sites in predicted presences for each of the 1000 permutations where their ^14^C dates were randomised. We then compared this number with the observed number of sites in predicted presences when assigned their real ^14^C dates.

### Inference of population structure and admixture events

We run the programme ADMIXTURE (26) on our pruned dataset. ADMIXTURE estimates for everyone the proportions of the genome originating from K ancestral populations, K being specified a priori. The programme was run at K values from 1-8 with CV error estimation and default values for fold iteration (v=5). We retained the results providing the lowest cross-validation (CV) error across iterations. In our case, this corresponded to K=3 (Fig. S12).

### Reconstruction of recent effective population sizes of CAHG and Bantu-speaking populations

Changes in effective population sizes (N_e_) in 9 CAHG populations as well as 15 Bantu-speaking farming populations from Cameroon and Gabon were traced over the period spanning 150-20 generations ago (4,500-600 years), after confirming through simulations that reliable N_e_ estimates could be obtained for this period given the structure of our dataset (Table S3; SM text, Fig. S14). *Ne* was estimated for each community using IBDNe software (22) on our non-pruned dataset after removing centromere and telomere regions where IBD density is particularly high (4,22). Data were initially phased with Beagle 5.1 using the haplotype maps available from the IBDNe website (http://bochet.gcc.biostat.washington.edu/beagle/genetic_maps/). Segments of IBD were identified using refinedIBD (86), after which they were merged and missing segments were filled with merge-ibd-segments from refinedIBD. Mean *Ne* values were calculated for each generation in each community, and generations were converted to calendar years before the present (BP) by assuming a generation time of 30 years, the recommended value for preindustrial societies in genetics-based studies of population divergence (87).

To control for the effect of admixture on the effective population sizes of CAHG, we then used RFMix to estimate local ancestry proportions along the genome in order to conduct ancestry-specific IBD analysis in those CAHG populations showing evidence of admixture (>10% median Bantu-associated ancestry according to the output of ADMIXTURE) (88)(Table S6). Following Perry et al. (67) and Ioannidis et al.(89), reference panels for each population were created by selecting unadmixed Bantu individuals (with >98% Bantu ancestry) according to the output from ADMIXTURE K=2, which illustrates the split between CAHG and Bantu populations (Fig. 2D). These were then paired with each CAHG population and ADMIXTURE was rerun on each pair to select appropriate reference panels for each specific case. Individuals with admixture proportions <10% were chosen for the reference panels in each separate population. RFMix v1 was run with the recommended two expectation maximization iterations, a 2-millimorgan window size and -n 5 as recommended in the manual for non-trio-phased datasets to determine local ancestry along the genome.

Phasing results from RFMix were then matched to phasing from Beagle and IBD haplotypes following the methods from Browning et al. (88) for ancestry-specific IBDNe (ASIBDNe). To account for uncertainty in ancestry estimation, we randomly assigned ancestry to each SNP according to a weighted estimation of ancestry per SNP (88). For example, a locus identified as 80% likely to belong to ancestry 1 and 20% likely to belong to ancestry 2 by RFMix would be randomly assigned to one of the two ancestries with ancestry 1 weighted at 0.8 and ancestry 2 weighted at 0.2. ASIBDNe was then used to calculate effective population size for the last 150 generations for each of the CAHG populations on each of the ancestry components separately.

### Inferring recent genetic connectivity between CAHG

IBD sharing between populations provides insights into recent contact or recent common ancestry. Since CAHG have a deep divergence(1), shared IBD blocks can be used to infer recent contact (39-40,89). We analyzed IBD blocks in three categories: 1–5, 5–10, and >10 cM.

We used refinedIBD (86) to identify shared IBD blocks between each pair of individuals in our dataset without centromeres and telomeres, and homozygous-by-descent blocks within each individual (40). Then, we merged IBD blocks within a 0.6-cM gap and allowed only one inconsistent genotype between the gap and block regions using the program merge-ibd-segments from BEAGLE utilities (88). These results were used to create three data sets based on the length of identified IBD blocks: 1–5 cM, 5–10 cM, over 10 cM. For each data set, we summarised IBD sharing between populations, by considering the probability that an individual selected at random from population A shared IBD segments of the specified length with an individual selected at random from population B (89). This was done, for each pair of populations, by dividing the total number of individual pairs connected by IBD segments of the length of the corresponding dataset by the total number of possible dyads for that pair of populations (90). We only kept the pairs with at least two shared blocks (4 for the range of 1–5 cM) to reduce noise and false positives. As a complementary measure of post-divergence gene flow between CAHG populations, we also calculated private allelic richness (per variable site) of alleles shared by pairwise combinations of the 9 CAHG populations using ADZE (91)(SM Text).

## Data availability

Data on the size and approximate location of camps are available as supplementary files (Data S1). Exact geographical coordinates remain confidential for protection of the privacy of the indigenous people involved.

Access to human genome-wide SNP data in the European Genome-Phenome Archive was granted via accession code EGAS00001002078 and EGAS00001000908.

Data on the location and dating of hunter-gatherer archaeological sites are available as supplementary files (Data S2).

Data on the location and dating of initial ^14^C date compilation are available as supplementary files (Data S3).

Data on the location and dating of hunter-gatherer archaeological sites after aggregating multiple dates within the same grid cell and 1000- or 2000-time interval are available as supplementary files (Data S4).

## Code availability

All code to reproduce the results reported in this paper will be made available upon acceptance of the manuscript.

## Acknowledgments

The authors wish to acknowledge all the anthropologists that have facilitated ethnographic data on hunter-gatherer camps and Etienne Patin for providing the genetic data. They also want to thank Matt Peeples for help in calculating the connectivity between hunter-gatherer camps, Luisa Espinós Iglesias for assistance with visualization and Bastiaan Star for support and guidance. Last, we are grateful to Hanna Kokko, and Inez Derkx for helpful discussions of the manuscript.

